# CUSP: Complex Spike Sorting from Multi-electrode Array Recordings with U-net Sequence-to-Sequence Prediction

**DOI:** 10.1101/2025.11.18.689109

**Authors:** Chenhao Bao, Robyn Mildren, Adam S. Charles, Kathleen E. Cullen

## Abstract

**Background:** Complex spikes (CSs) in cerebellar Purkinje cells convey unique signals complementary to Simple spike (SS) action potentials, but are infrequent and variable in waveform. Their variability and low spike counts, combined with recording artifacts such as electrode drift, make automated detection challenging.

**New Method:** We introduce CUSP (CS sorting via U-net Sequence Prediction), a fully automated deep learning framework for CS sorting in high-density multi-electrode array recordings. CUSP uses a U-Net architecture with hybrid self-attention inception blocks to integrate local field potential and action potential signals and outputs CS event probabilities in a sequence-to-sequence manner. Detected events are clustered and paired with concurrently detected SSs to reconstruct the complete Purkinje cell activity.

**Results:** Trained on cerebellar neuropixels recordings in rhesus macaques, CUSP achieves human-expert performance (F1 = 0.83 ± 0.03) and even captures valid CS events overlooked during manual annotation.

**Comparison with Existing Methods:** CUSP outperforms traditional and state-of-the-art CS and SS sorting algorithms on CS detection. It remains robust to waveform variability, spikelet composition, and electrode drift, enabling accurate CS tracking in long-term recordings. In contrast, existing methods often show false-positive biases or degrade under drift.

**Conclusions:** CUSP provides a scalable, robust framework for analyzing burst-like or dynamically complex spike patterns. Its generalizability makes it valuable for large-scale cerebellar datasets and other neural systems, such as hippocampal pyramidal cells, where complex bursts are critical for computation. By combining expert-level accuracy with automation, CUSP offers a broadly applicable solution for studying information coding across circuits.

## 1. Introduction

The cerebellum plays a critical role in coordinating movement and ensuring that actions are precisely timed. By comparing intended motor commands with actual outcomes, it uses sensory feedback to generate rapid corrections and support ongoing motor learning. When cerebellar function is impaired, movement initiation is preserved, but control becomes compromised, resulting in unsteady, dysmetric actions and difficulties with balance, posture, and adaptation (Selhorst et al., 1976; Bakker et al., 2006; Horak & Diener, 1994; Bastian & Thatch, 1995; Morton & Bastian, 2006; Smith & Shadmehr, 2005; Maschke et al., 2004; Sokolov et al., 2017). A central challenge in understanding cerebellar computations lies in measuring the activity of Purkinje cells (PCs)—the sole output neurons of the cerebellar cortex. PCs produce two distinct types of spikes: high-frequency simple spikes (SSs), driven by mossy fiber input via granule cells, and low-frequency complex spikes (CSs), generated by climbing fiber input from the inferior olive (Eccles et al., 1966; Thach, 1967). CSs occur at ∼1 Hz and are characterized by a large initial spike followed by a variable number of smaller spikelets (Llinás & Sugimori, 1980; Davie et al., 2008). Unlike the stereotyped shape of SSs, CSs vary substantially across cells, sessions, and even over time within a recording (Yang & Lisberger, 2014; Lang et al., 2014; Zang et al., 2018; Burroughs et al., 2020; Markanday et al., 2020; Sedaghat-Nejad et al., 2021). Such variability stems from multiple anatomical and physiological factors—including the branching patterns of PC dendrites, climbing fiber input dynamics, voltage state, instantaneous climbing fibre firing rate, and influence of molecular layer interneurons (e.g., Zang et al., 2018; Rowan et al., 2018). Consequently, CS events differ in spikelet number, timing, and amplitude, even within the same neuron, making the use of standard spike sorting techniques to detect CSs in extracellular recordings difficult, particularly in high-density datasets.

Despite this challenge, accurate detection of CSs is essential for probing cerebellar learning mechanisms. Theoretical models of cerebellar function—most notably the Marr-Albus-Ito hypothesis—propose that CSs convey motor error signals that instruct synaptic plasticity (Marr, 1969; Albus, 1971; Ito, 1972). While considerable evidence supports this framework (e.g., Medina & Lisberger, 2008; Herzfeld et al., 2015; Kostadinov & Häusser, 2022), recent findings have raised questions about how and when CSs encode behaviorally relevant signals (Catz et al., 2005; Dash et al., 2010; Junker et al., 2018; Kostadinov et al., 2019; Ohmae & Medina, 2015; Streng et al., 2017). Resolving these debates—and linking CS activity to motor learning—requires detecting CSs with high fidelity across large-scale recordings.

Extracellular electrophysiology offers millisecond-resolution access to PC output, and high-density multi-electrode arrays (MEAs) have expanded the scope of these recordings across cerebellar layers and lobules. However, spike sorting from MEA data remains difficult (Koukuntla et al. 2025), particularly for CSs. Most traditional algorithms are optimized for detecting SSs (Bao & Charles 2023), and perform poorly on the sparse, variable, and complex nature of CS waveforms. Early methods leveraged statistical features such as the post-CS pause in SS firing (Eccles et al., 1966; Bell & Grimm, 1969). More recent methods have adopted waveform- and LFP-based strategies, including PCA of low-frequency content (Zur & Joshua, 2019), semi-automated clustering with human supervision (Sedaghat-Nejad et al., 2021), and deep learning applied to single-channel signals (Markanday et al., 2020). While these tools represent important advances, they typically operate on single electrodes and thus fail to leverage the rich spatial information now available from high-density MEAs. As a result, they remain vulnerable to waveform variability, electrode drift, and noise—persistent challenges in naturalistic and long-term recordings. These limitations underscore the need for a new method designed to exploit spatial structure to improve accuracy, robustness, and scalability.

To overcome these limitations, we developed CUSP—Complex spike sorting via U-net Sequence Prediction—a fully automated deep learning framework for identifying CSs from MEA recordings. CUSP casts spike sorting as a sequence-to-sequence prediction task and incorporates hybrid self-attention inception blocks into a U-net architecture, allowing it to integrate LFP and AP signals across space and time. The model outputs probabilistic CS detections, which are then clustered and paired with concurrently detected SSs to reconstruct the full Purkinje cell firing profile. Evaluated on cerebellar recordings from rhesus macaques, CUSP achieves expert-level performance and outperforms traditional and state-of-the-art methods. By enabling robust, scalable, and fully automated CS sorting, and by remaining robust to waveform variability and electrode drift, the framework has the potential to be more broadly applicable to other neural systems, such as the Hippocampus, where burst-like or dynamically complex spike patterns pose similar challenges.

## 2. Methods

As Purkinje cells are defined by both CS and SS events (**Fig. 1**), our complex spike sorting runs in parallel to canonical spike sorting and is subsequently merged with the SS sorting results. Specifically, we first preprocess the raw MEA recordings and then input into our ML model. The model identifies CS events by producing an indicator time series that represents the probability of a CS event at each sample time. Contiguous high-probability time-points in this series then allow us to identify each CS candidate, which is then used for downstream clustering of CS events into individual units. Finally, the output clusters are paired with the SS sorting results to produce full representations of putative Purkinje cell activity (**Fig. 2**).

**Fig 1.**
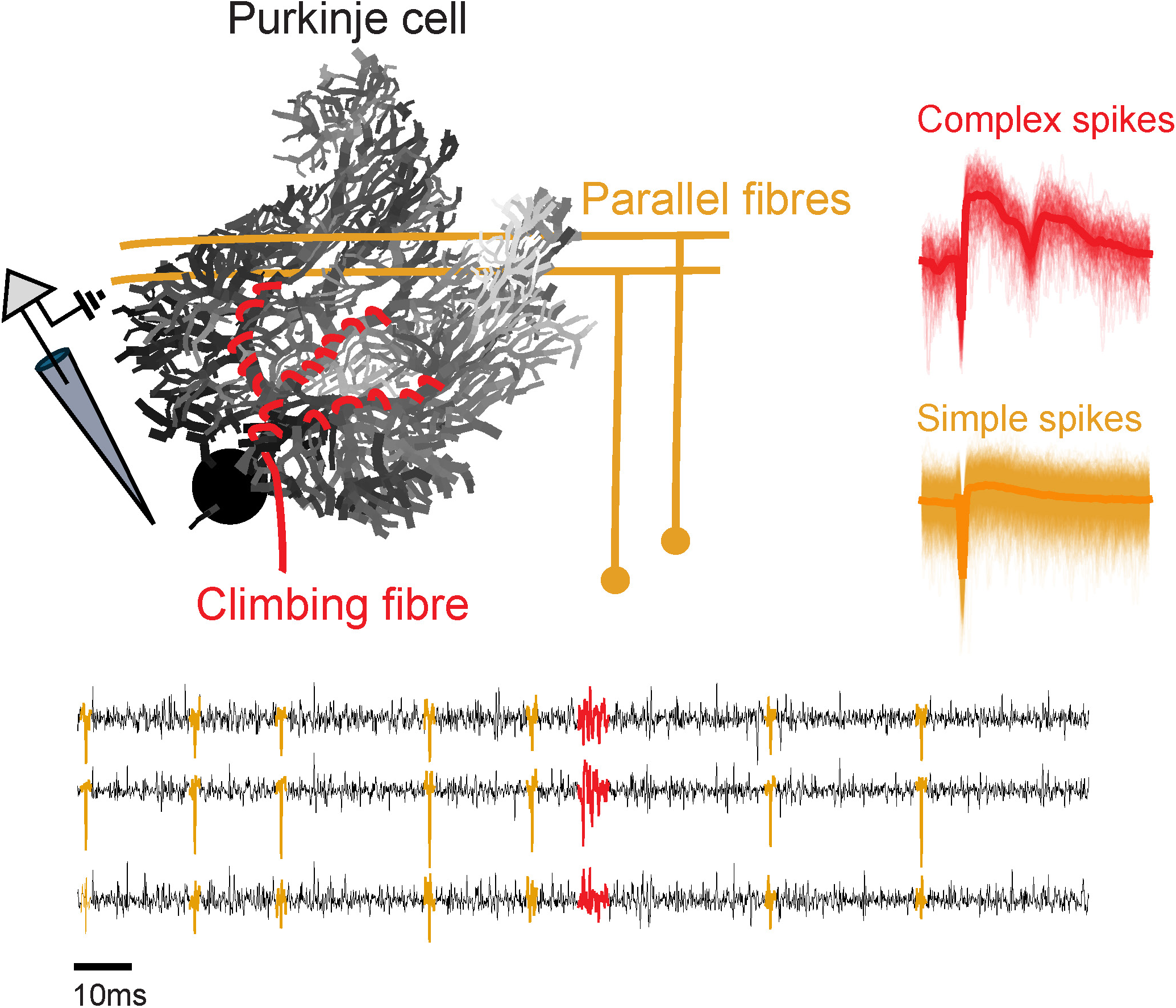
Illustration of Purkinje cell simple and complex spikes. **(Top)** In the cerebellar cortex, Purkinje cells generate simple spikes (SS, orange) from parallel fiber input and complex spikes (CS, red) from climbing fiber input. **(Bottom)** Example extracellular recording. Time series traces (black) and overlaid SS (orange) and CS (red) waveform templates. SS have a stereotypical depolarization–repolarization cycle, whereas CS feature multiple spikelets following the initial depolarization.

**Fig 2.**
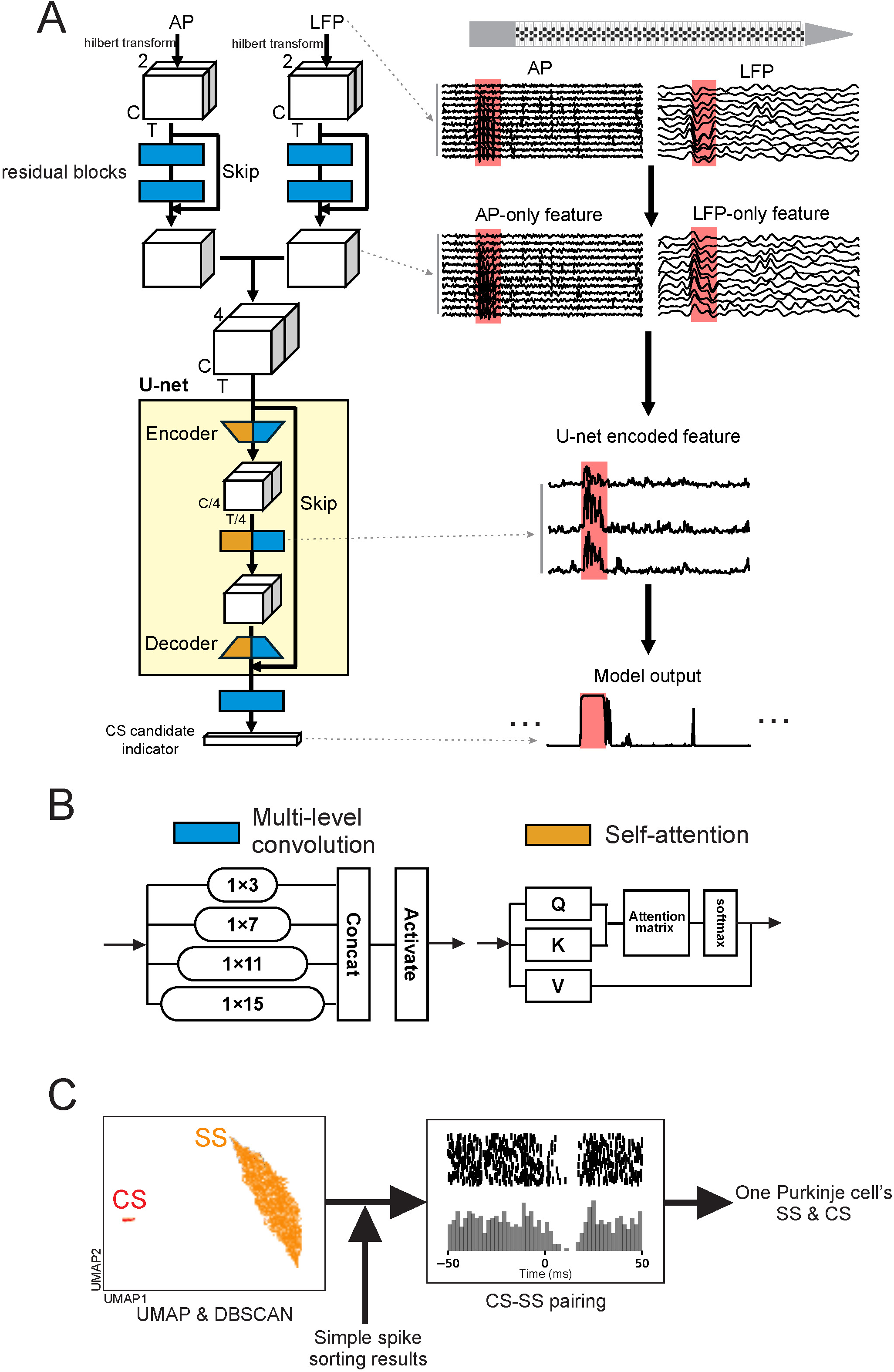
Schematic of our Purkinje cell spike sorting pipeline. **(A)** CUSP model architecture (left), and example feature visualization at each stage of the model (right). Multi-electrode array (MEA) recordings are preprocessed and filtered to the action potential (AP; 300-3000 Hz) and local field potential (LFP; 30-400 Hz) bands. Example AP and LFP bands are shown across a subset of channels; red boxes indicate expert-labeled CS events. Hilbert-transformed AP and LFP signals are input to the residual blocks to obtain LFP-AP-specific features. These features are stacked and sent into a U-net with hybrid self-attention inception blocks to extract combined LFP/AP features that are used to predict CS events through a sequence-to-sequence network. The model outputs a CS candidate indicator sequence, where higher values indicate a greater probability of a CS event. **(B)** Schematics of the multi-level convolution and self-attention modules in the hybrid self-attention inception block. **(C)** Postprocessing steps identify a Purkinje cell embedding the identified CS waveforms via uniform manifold approximation and projection (UMAP) and clustering via density-based spatial clustering of applications with noise (DBSCAN). These clusters are then combined with SS sorting results by matching the CS-SS pairs based on their cross-correlogram (CCG), where SS events are, as in the depicted histograms, expected to decrease following a CS event.

### 2.1 Deep neural network based CS sorting

#### Data collection and preprocessing

Our CS sorting model was trained and evaluated on a subset of extracellular recordings from the cerebellar cortex (specifically the nodulus/uvula in the posterior vermis) of two rhesus macaques from previous study (Mildren et al., 2025). Neural activity was recorded using 128-channel silicon read-write electrodes (IMEC), where the electrodes are arranged in a zig-zag pattern within a ∼1.3 mm length and 0.06 mm width recording region. This electrode spacing, density, and impedances are comparable the Neuropixels-NHP electrodes. Recorded signals were amplified, bandpass filtered (0.1-7500 Hz), digitized, and streamed at 30 kHz from a set of data acquisition systems (four 32-channel RHS stim/recording head stages and an RHS stim/recording controller, Intan technologies).

The raw multi-channel signals from the data acquisition system were preprocessed before being sorted. First, the DC component within each channel was removed to center each time-trace. Next, all channel signals were re-referenced to a common median value. Each channel signal was then further re-referenced to a local 12-channel common median value, as the common median referencing has been previously shown to improve signal to noise ratio for spike detection relative to the common average referencing (Rolston et al., 2009). Finally, the high and low frequency portions of the channel were separated to better enable the subsequent ML algorithm to leverage the distinct features in the local field potential (LFP) and spiking action potential (AP) portions of the voltage traces.

To isolate the LFP and AP bands, each channel signal was filtered to the corresponding frequency ranges (30-400 Hz for LFP and 300-3000 Hz for AP) via zero-phase 4th-order Butterworth bandpass filters. We note that in theory, a complex enough ML model could identify high and low-pass features from the raw time series. However, we found that the model trains more efficiently and effectively when the two bands are separated and used as separate input streams, which allows the ML model to focus on identifying features within the LFP and AP bands that are known to carry complementary information about CS events (Perelman & Ginosar, 2007).

### ML model

Our CS sorting model, CUSP (schematized in **Fig. 2A, left panel**), accepts input sequences in a sliding window manner, with each sliding window segment containing a 50-ms time length (1500 samples with a 25-ms step) and 12-channel local LFP and AP signals (example traces shown in **Fig. 2A**, right upper panel), for a total of 36000 inputs across the 24 input time series. We first perform a Hilbert transform on the LFP and AP segments to acquire their real and imaginary components, which contain both amplitude and phase information, and then use residual blocks (He et al., 2016) with multi-level convolutional layers (i.e., inception block) (Szegedy et al., 2015). Each inception block contains four scales of convolutional kernels with the reception field of 1×3, 1×7, 1×11, and 1×15 (blue block, **Fig. 2B**). The extracted LFP-only and AP-only features derived from the residual blocks are then stacked and sent into our neural network with a U-net structure.

The U-net structure forms an information bottleneck to compress the useful information by building two symmetrical paths---the encoder and decoder---joined with skip connections. The encoder performs both sample-wise and channel-wise downsampling, progressively condensing the feature information while the decoder reconstructs the fine-grained information back by upsampling, and recovers precise localization via the skip connections from the original (Ronneberger et al., 2015). Within our U-net structure, we build each block with a hybrid self-attention inception, with the multi-level convolution and self-attention modules (Vaswani et al., 2017) (orange block, **Fig. 2B**) running in parallel.

The hybrid self-attention inception block allows CUSP to capture both the global dependencies from self-attention as well as the local pattern hierarchies from the multi-level convolution, and therefore leads to a more robust and comprehensive representation of CS features. The final output from CUSP is a CS candidate indicator, where a higher value indicates a higher probability of a CS event (**Fig. 2A**, right bottom panel). By producing a scalar value rather than a binary output, we can better quantify the CS sorting model confidence and control the final classification through flexible thresholding. CUSP thus represents fully sequence-to-sequence processing, with the exact sample timing preserved in the output to best identify the onset and duration of each CS event.

### Model training, validation, and inference

We trained the CUSP network on the macaque cerebellar cortex dataset with manual labeling of complex spikes. To define ground truth for supervised training of the model, three experts independently labeled the timing of CS, and the CS labels mutually agreed among experts were selected as the ground truth. Several data augmentation techniques were used to enrich our training dataset as well as improve our CS sorting model’s robustness. Specifically, the data augmentation included channel drift, sample jitter, and voltage reversal, the latter of which was included to prevent overfitting on spike waveform peak/valley values.

CUSP’s ML cost function is a binary cross-entropy loss with a logistic function, which we optimized with the Adam stochastic optimization (Kingma et al., 2014). We coded CUSP’s ML block in PyTorch (Paszke et al., 2019), and performed all training, testing, and analyses on a standard high-performance computing machine with NVIDIA Tesla A100 GPU. We trained CUSP for approximately 1000 epochs with a batch size of 96 on our complex spike dataset. The total training time was approximately 72 hours on a single NVIDIA A100 GPU. The trained CUSP was then applied to neural recording data to detect CS in an offline processing manner, and we calculated the inference speed on a single NVIDIA GeForce RTX 4050 Laptop GPU, with an average inference time cost 0.429±0.011 second per 1-second recording data.

### 2.2 Postprocessing pipeline for determining one Purkinje cell

As CUSP’s ML block outputs CS estimates of the probability of an event, we post-process the output time series to detect each CS onset time. Specifically, we apply a threshold to the output trace, where the default threshold is chosen as the quantile value of 0.7 (approximately the mean + 1 standard deviation for the normal distribution, i.e., 0.683). We visualize the 2-D embedding results of the CS candidate based on their multi-channel waveforms using a nonlinear dimension reduction method, uniform manifold approximation and projection (UMAP) (McInnes et al., 2018), and find the CS clusters via density-based spatial clustering of applications with noise (DBSCAN) (Ester et al., 1996). We introduce flattened auto-convolution features of each identified CS candidate multi-channel waveform for clustering, which improves the clustering robustness against noise.

After identifying the CS clusters, we combine the CS information with the SS sorting results to find the corresponding SS activity for each CS cluster. The CS-SS pair is identified by checking whether the CS-SS cross-correlogram (CCG) presents a strong SS firing inhibition after CS onset (**Fig. 2C**). Each CS-SS pair identified in this way defines the complete neural activity of one putative Purkinje cell.

### 2.3 Evaluation and benchmarking

We performed leave-one-out cross-validation (LOOCV) to evaluate our CS sorting model, leaving the MEA recording data of one Purkinje cell out for validation and using the MEA recording data of the rest of the Purkinje cells for training. Each recording session was conducted independently over a period spanning several months.

Following each session, the complex spike activity of each individual Purkinje cell was manually annotated by human experts. Consequently, the electrode recording site and the surrounding neuronal population differed across sessions, ensuring that recordings were obtained from distinct cells and locations.

This procedure ensured that no information leakage occurred between the training and evaluation datasets when separating one Purkinje cell recording session from the rest of the dataset. We applied leave-one-out cross-validation (LOOCV) across a total of 17 recording sessions, each containing human expert–labeled Purkinje cell complex spike activity. In each iteration, data from 16 Purkinje cells were used for training and data from the remaining Purkinje cell were reserved for validation. This process was repeated 17 times so that each Purkinje cell served once as the validation set. On average, excluding one Purkinje cell removed 77.4 CS event instances from the training data (see **Fig. S1**). Each detected CS cluster was compared with the ground truth CS labels in a binary hit-or-miss manner, defined as whether the detected CS presents within a 5ms tolerance window of the ground truth CS. This spike timing measurement examines how well the CS sorting model captures the precise CS timing information and is critical for further testing temporal encoding and firing rate calculation (Bao & Charles, 2023).

We constructed a fair expert performance comparison framework between our CUSP and an individual human expert to examine whether our CS sorting model is aligned with human expert manual CS sorting performance. We used two of three experts’ mutually agreed CS labels as the human expert consensus ground truth to evaluate both our CS sorting model performance and the third individual human expert’s performance. We repeated this process three times, alternating between the three experts, in a round-robin manner. We calculated the precision, recall, and F1 scores for both our CS sorting model and individual human expert manual sorting:

Precision = (# matched CS)/(# detected CS),

Recall = (# matched CS)/(# ground truth CS),

F1 = (2*Precision*Recall)/(Precision+Recall).

In addition, we built a unified benchmarking framework to evaluate our CS sorting model against other existing CS sorting methods (Zur & Joshua, 2019; Markanday et al., 2020), as well as some representative general propose spike sorting methods (Pachitariu et al., 2024; Yger et al., 2018; Garcia & Pouzat, 2015; Chung et al., 2017) implemented through a unified API framework called SpikeInterface (Buccino et al., 2020). We used the consensus annotations across all three expert-agreed CS labels as the ground truth. The 3-annotator consensus serves as the most trustworthy ground truth in the context of minimizing human biases (Wilson et al., 2003; Scheuer et al., 2017) and thus provides the most faithful characterization of each algorithm’s intrinsic systematic biases. Similarly, the precision, recall, and F1 scores for all algorithms were calculated and compared.

## 3. Results

### 3.1 Our complex spike sorting method reaches human expert consensus level without individual bias

To establish a robust baseline for comparing CS sorting algorithms, we first examined our ground truth CS labels from human experts. Among the three experts’ independent complex spike labels, we found a gap between individual expert and multiple expert consensus CS labeling results, with the percent of CS events mutually agreed upon by any two different experts being only 73.7±3.6% (mean ± standard error, same below). The consensus level between three experts further dropped to 68.8±3.8% across all Purkinje cells in our dataset (**Fig. 3A**). These numbers suggest that although most CSs can be agreed upon by human experts, there exists a significant partition of CSs that failed to be consistently recognized by human annotators. The failure to consistently label so many CS events likely comes from the high variability in CS waveforms due to effects such as noise, electrode drift, variable spikelets, as well as the variability in individual human expert biases during their manual CS labeling.

**Fig 3.**
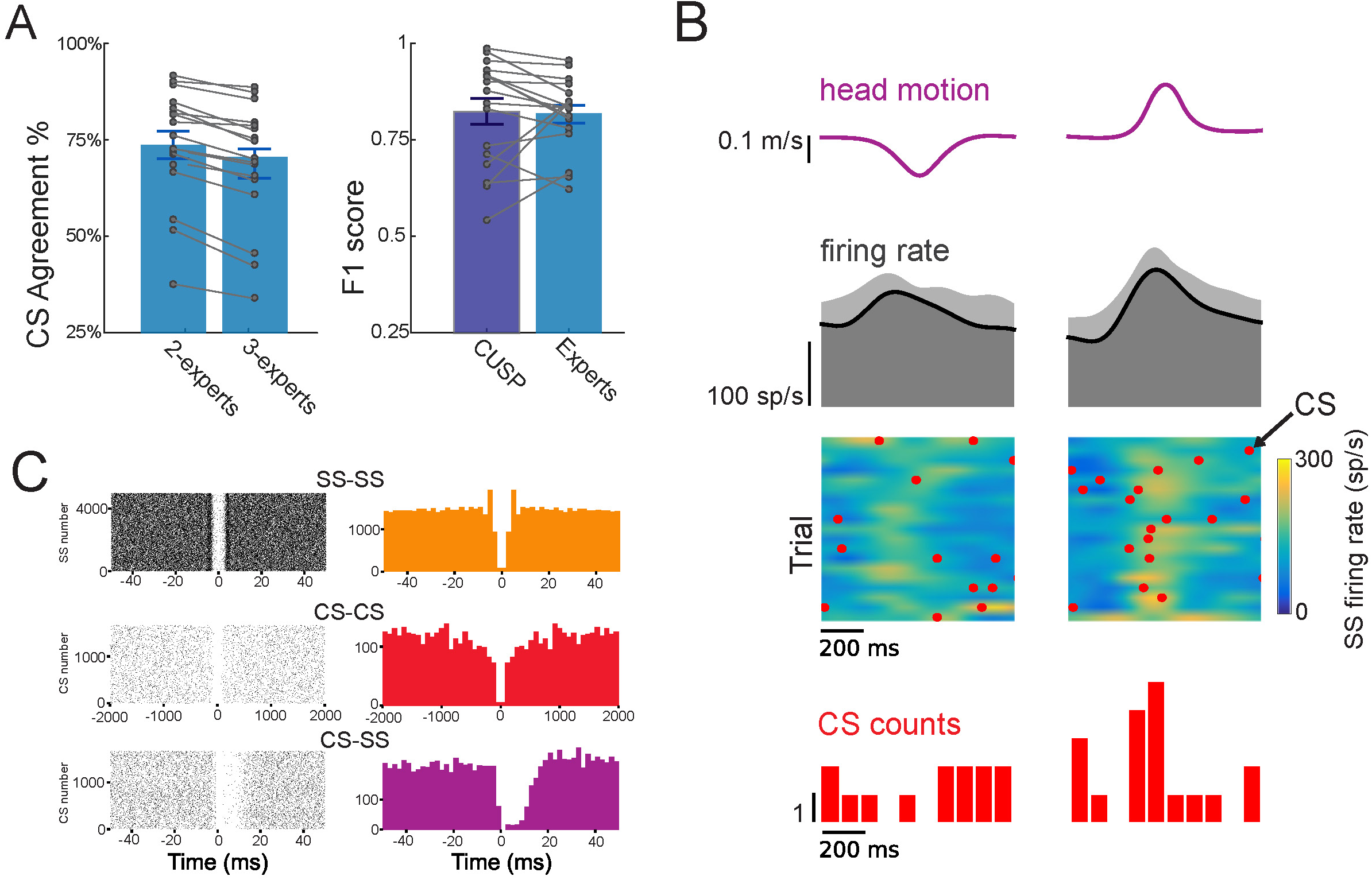
Complex spike sorting results from CUSP. **(A)** Mutual CS agreement between two human experts compared to three experts (left) and F1 score between CUSP and human experts (right). CUSP shows CS sorting performance equivalent to cross-expert consensus (human expert F1 = 0.82±0.02, CUSP F1 = 0.82±0.03, p=0.76). Each dot represents the performance for an individual Purkinje cell and error bars represent standard error. **(B)** Example CUSP results on CS responses during sensory stimulation evoked by head translations in the posterior (negative) and anterior (positive) directions. Head velocity (top), mean SS firing rate (2^nd^ row) and heat plot (3^rd^ row) demonstrates SS firing rate across trials. Red dots indicate CS events and the red histogram (bottom) depicts the probability of CS firing during stimulation, where there was a high probability of CS events aligned with anterior head motion. **(C)** Auto-correlograms for CS (top) and SS (middle), and cross-correlograms between CS and SS for the Purkinje cell in panel B.

We next evaluated our algorithm’s performance against typical human expert annotations. Specifically, we iterated through all pairs of expert annotators, and for each pair, we used the consensus between those annotators as ground truth correct CS events. We then compared the F1 score between our CS sorting algorithm and the third, remaining human expert, against the 2-annotator consensus ground truth (see Methods). We computed the final accuracy numbers as the mean and standard deviation across all expert pairs. This procedure enabled us to 1) compare the relative performance of each algorithm to expert performance, and 2) remove the potential bias of any particular pair of experts. We found that individual human expert performance had an F1 score of 0.82±0.02 (**Fig. 3B**, red), while our CS sorting model achieved a comparable F1 score of 0.82±0.03 F1 (**Fig. 3B**, blue), which was not significantly different from the human expert performance (p=0.76). This comparison demonstrates that our CS sorting model performs at a level equivalent to that of typical human expert manual labeling.

### 3.2 CUSP out-performs current methods on the benchmarking CS sorting datasets

Given that our CS sorting model achieved expert-level labeling performance, we further evaluated against the current state-of-the-art CS sorting algorithms using a unified benchmarking framework, where we used the consensus annotations across all three expert-agreed CS labels, which minimize human biases and faithfully reflect each algorithm’s intrinsic systematic biases.

As above, we compared performance on our Purkinje cell complex spike dataset, with the full set of comparison algorithms spanning both single-channel and multi-channel sorters and including:

1. Canonical single channel LFP/AP voltage thresholding, with PCA/UMAP embedding and DBSCAN clustering;
2. (Markanday et al., 2020): A single-channel CS detection model using a shallow convolutional neural network, translated from the architecture of model initially developed to detect saccadic eye movement (Bellet et al., 2019);
3. Spyking Circus2 (Yger et al., 2018): A multi-channel general-purpose spike sorting algorithm based on template matching and Density Peak Clustering;
4. Tridesclous2 (Garcia & Pouzat, 2015): A multi-channel general-purpose spike sorting algorithm based on template matching and a customized clustering;
5. Mountainsort5 (Chung et al., 2017): A multi-channel general-purpose spike sorting algorithm based on a non-parametric ISO-SPLIT clustering;
6. Kilosort4 (Pachitariu et al., 2024): A multi-channel general-purpose spike sorting algorithm based on template matching and graph clustering

We computed the F1 score performance of each algorithm on our benchmarking dataset (**Fig. 4**, pink). Notably, there is an obvious performance gap between single-channel sorting algorithms and multi-channel sorting algorithms, where the multi-channel sorting algorithms always hold better F1 scores than single-channel sorting algorithms. Prevalent general-purpose spike sorting algorithms, including Kilosort4 (Pachitariu et al., 2024), Spyking Circus2 (Yger et al., 2018), Tridesclous2 (Garcia & Pouzat, 2015), and Mountainsort5 (Chung et al., 2017), all of which use multi-channel recording signals as inputs to retrieve spike spatial feature information and determine each unit’s nearest channel position. In the context of CS sorting, leveraging spatial information is even more necessary for accurate CS detection (F1 score 0.64±0.09, 0.68±0.06, 0.75±0.06, and 0.78±0.07, for Kilosort4, SpykingCircus2, Tridesclous2, and Mountainsort5, respectively). Our CS sorting model, which also uses the full multi-channel recording signal as inputs, presented the highest 0.83±0.03 F1 score in the benchmarking dataset.

**Fig 4.**
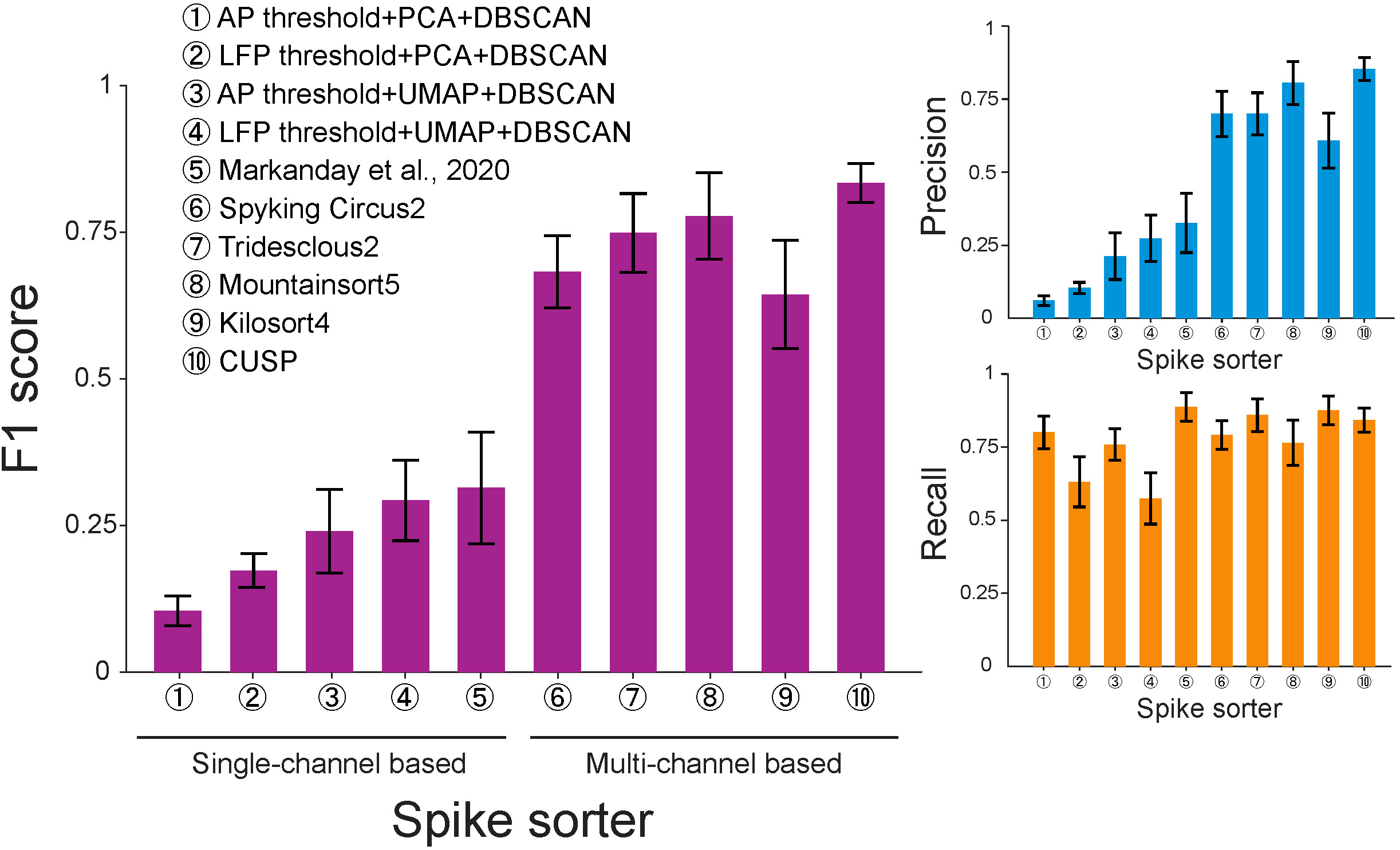
Benchmarking the performance of different spike sorting methods on our multi-electrode array recordings from Purkinje cells in the cerebellar cortex. **(Left)** Leave-one-out cross-validation F1 score of CUSP compared to four variants of single-channel thresholding, a previously published single-channel complex spike sorting algorithm from Markanday et al., 2020, and four recent and/or widely used sorting algorithms: Kilosort4, Spyking Circus2, Tridesclous2, and Mountainsort5. CUSP achieved the highest F1 score of 0.83±0.03. **(Right)** Same histogram for precision (top) and recall (bottom), showing the source of errors driving the F1 score. While Recall was relatively high across multielectrode sorters, precision was highest with CUSP reflecting the best accuracy for positive predictions.

To determine whether CUSP achieved significantly higher performance than existing algorithms, we preformed a paired permutation test with 100,000 random repetitions, applying reweighting based on the total number of expert-labeled complex spikes to compensate for differences in session duration and CS firing rates. The results showed that CUSP’s benchmarking performance was statistically higher than that of the state-of-the-art spike sorter (Mountainsort5) across all metrics including F1 score (0.83±0.03 vs 0.78±0.07, p<1.00e-5), precision (0.85±0.04 vs 0.80±0.07, p=2.00e-5) and recall (0.84±0.04 vs 0.76±0.08, p<1.00e-5) (**Fig. S3**).

As opposed to multi-channel methods, the single-channel sorting algorithms show a systematic disadvantage due to limited access to spatial information surrounding each single channel, and therefore struggle in identifying the correct CS waveforms from the background of environmental noise and crowded multi-unit activities. The canonical thresholding method based on single-channel AP/LFP signals only presents minimal CS sorting improvement when using PCA for dimensional reduction and DBSCAN for clustering (F1 score 0.10±0.02 and 0.17±0.03, respectively). By switching from linear PCA embedding to nonlinear UMAP embedding, both the canonical AP/LFP thresholding only show slightly increased F1 scores (0.24±0.07 and 0.29±0.07, respectively). Even the existing single-channel CS detection model (Markanday et al., 2020) that we reproduced on our dataset was still not sensitive enough for isolating CSs (F1 score 0.31±0.09).

We next investigated the detailed precision (**Fig. 4**, orange) and recall scores (**Fig. 4**, blue) for single-channel sorting algorithms and multi-channel sorting algorithms to investigate whether type I error (false positives) or type II error (false negatives) is limiting the performance increase from single-channel sorting algorithms to multi-channel sorting algorithms. We found that while some increases in recall scores were shown, there were even larger increases in precision scores when shifting from single-channel sorting algorithms to multi-channel sorting algorithms, which led to the observed increases in F1 score. This suggests that the algorithms based on the single-channel signal seem to always be biased towards type I error (false positives), generating an overwhelming amount of hallucinated CS detections. On the other hand, algorithms based on the multi-channel signals seem to leverage the spatial feature conveyed within the inter-channel information to help themselves more precisely identify the correct CS waveforms and get rid of unconvincing detection results, and their higher F1 scores are mainly attributed to improved precision scores.

### 3.3 CUSP is robust to spikelet variability and external perturbations

Finally, we investigated how well CUSP performs in response to challenging, but common, confounding effects that increase the CS waveform variability. First, we examined the robustness of our model to intra-Purkinje-cell variability by evaluating our model on those Purkinje cells with changeable CS waveforms in response to different climbing fiber inputs, based on the human expert annotations. We found that despite variable spikelet numbers and inter-spikelet intervals, our CS sorting model successfully identified multiple complex spike waveform variants (**Fig. 5A**).

**Fig 5.**
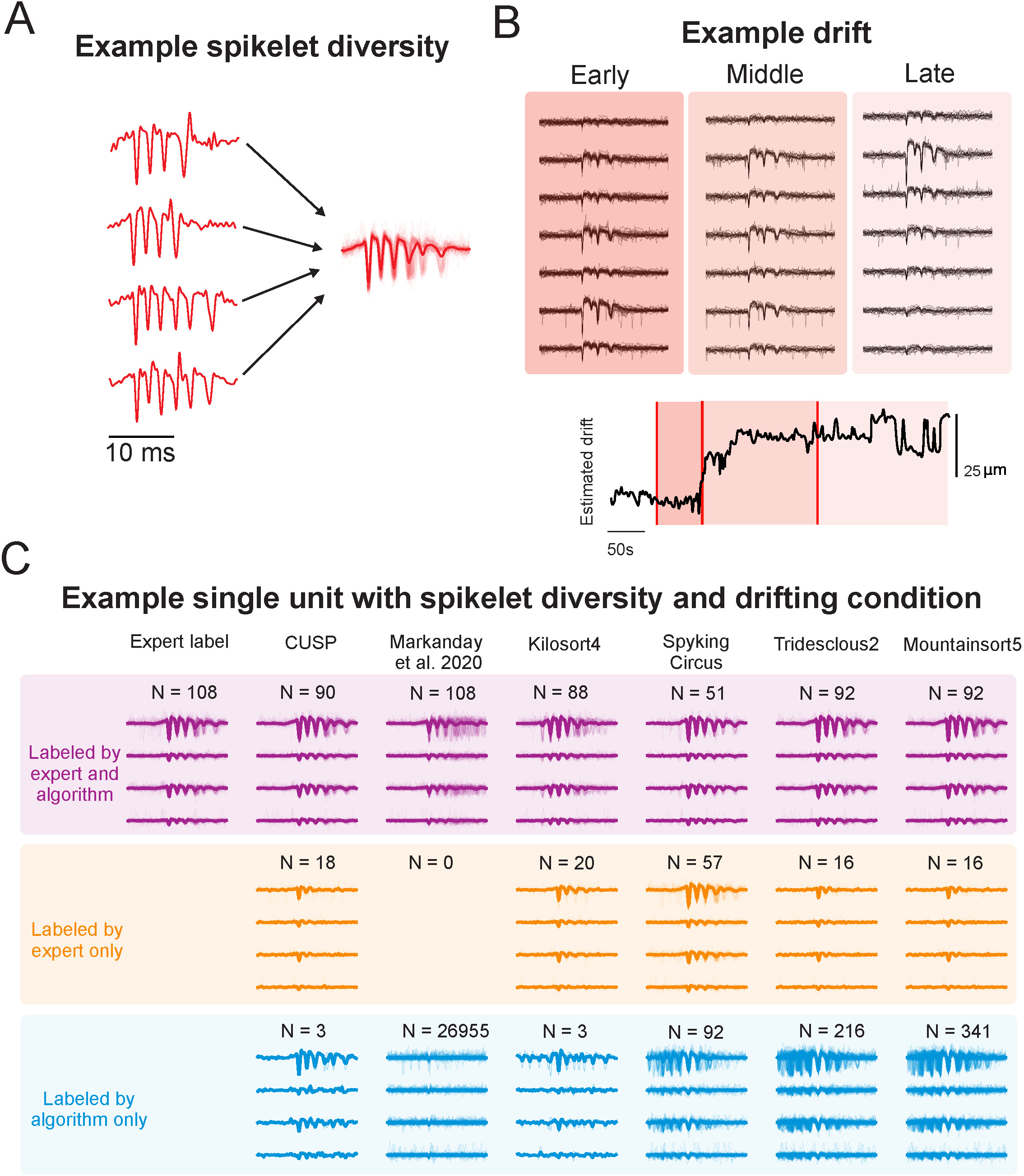
CUSP’s robustness against waveform variability and drift leads to high performance in minimizing spike sorting errors. **(A)** CUSP can detect complex spikes from the same Purkinje cell despite substantial waveform variability. Individual red traces represent four example CS waveforms manually identified as from the same Purkinje cell despite substantial variation in the number of spikelets. **(B)** CUSP can also trace the same Purkinje cell’s complex spikes across channels during electrode drift. CS waveforms are shown across seven electrode channels during early, middle and late epochs of the recording. The estimated drift (bottom) highlights the presence of drift. **(C)** Example complex spike waveforms under spikelet diversity and drifting conditions identified using each complex spike sorter (columns of the plot). The purple “labeled by expert and algorithm” cluster shows the true-positive complex spikes that were detected by both the spike sorter and expert labeler. The yellow “labeled by expert only” cluster shows the waveforms that were identified by expert labelers but not the column’s corresponding algorithm. Finally, the blue “labeled by algorithm only” cluster shows waveforms that were labelled by the algorithm but not experts. This demonstrates CUSP likely correctly discards expert mislabels (N = 18, yellow) and finds additional CSs that were missed by experts (N = 3, blue).

To quantify algorithmic performance under these spikelet diversity conditions, we computed two complementary metrics designed to capture double-counting behavior. This analysis was motivated by the observation that highly variable spikelet waveforms from a single Purkinje cell can produce multiple time-shifted template-projections of the same complex spike, leading an algorithm to erroneously count a single event multiple times. Specifically, we quantified 1) the number of algorithm detections associated with each complex spike event – defined as the double-counting number (**Fig. S4A**), and 2) the temporal misalignment between each detected event and the expert-labeled CS onset time—defined as the double-counting time misalignment (**Fig. S4B**). Markanday et al., 2020, Spyking Circus, Tridesclous2, and Mountainsort5 all exhibited higher double-counting numbers for each CS, with misalignments broadly distributed before and after the CS onset, indicating that these algorithms frequently misclassified variable spikelets as independent spike events and thus generated excessive false-positives. By contrast, both CUSP and Kilosort4 showed near to one-to-one correspondence with expert labeled CSs; however, CUSP maintained a close-to-zero time misalignment relative to CS onset, whereas Kilosort4 exhibited broader post-onset misalignments and consequently failed to achieve precise onset detection. Thus, overall, CUSP demonstrated superior accuracy compared with all other algorithms in detecting CSs under spikelet diversity conditions.

We also tested our CS sorting model on MEA recordings that presented significant electrode drift to assess its stability in tracking drifted CS waveforms. We found CUSP successfully tracked the same complex spike that drifted across four channels (∼50µm) of the electrode from early to late stages of the recording (**Fig. 5B**). Importantly, if the algorithm relied on information from a single channel, tracking would not be successful since the peak amplitude of the CS waveform diminished until it was indistinguishable from noise on the channel where it was originally the largest. However, CUSP’s end-to-end utilization of multi-channel data allowed the CS to be identified from neighboring channels.

To identify further areas where our approach may provide robustness that enables improved performance over the existing state-of-the-art spike sorting algorithms, we explored specific instances representing discrepancies between the human expert labels and the algorithm’s results, such as instances involving complex combinations of spikelet diversity and electrode drift (**Fig. 5C**). CS where the manual and algorithm annotations matched were typically clearly visible against the background noise (purple) while unmatched expert labels (expert-exclusive; orange), were characterized by less prominent CS waveforms. In contrast, the algorithm-exclusive results (blue) showed CSs that had clear and intact CS waveforms, although they were missed by human experts, presumably due to annotator-specific biases or labeling fatigue. Existing sorting methods showed types of errors in these instances that drive their overall worse performance. Some methods detected false-positive CS waveforms that were misaligned with the ground truth CSs (e.g., matched results from Kilosort4); some showed false-negative biases in some situations and missed a large portion of good CSs (e.g., expert-exclusive results from Spyking Circus2); and most of the existing algorithms were biased towards overwhelming false-positives, specifically confused noise and/or spikelets (e.g., algorithm-exclusive results from Markanday et al., 2020, Kilosort4, Spyking Circus2, Tridesclous2, and Mountainsort5).

To better visualize the performance of each algorithm under conditions of spikelet diversity and electrode drift, we used UMAP to fit the embedded manifold from expert labeled CS waveforms and then transformed each algorithm detected CS waveforms onto the same embedded manifold (**Fig. S5**). The embedded results revealed that expert labeled CS waveforms formed multiple clusters due to waveform changes associated with drift and spikelet diversity. CUSP detected CS were closely aligned with expert labeled clusters, excluding less prominent CS-like waveforms. In contrast, Markanday et al. 2020, Spyking Circus, Tridesclous2, and Mountainsort5 all showed numerous false-positive detections that laid outside of the expert labeled CS waveform distributions. Kilosort4’s performance was slightly worse than CUSP with a greater proportion of detected CSs located near neighboring false-positive clusters and only marginally overlapping with expert-labeled CSs. Overall, CUSP produced the most robust and accurate results under these challenging conditions, consistent with the superior performance observed in our quantitative benchmarking.

We next evaluated whether CUSP’s performance generalizes to recording datasets beyond those used for expert-labeled training. Specifically, we applied the well trained CUSP model to additional recording sessions collected from different rhesus macaque subjects and across multiple cerebellar regions. Using this model, we successfully identified numerous well-defined CSs from our larger recording database (see examples of the CUSP-detected CSs in **Fig. S2**). These results demonstrate that CUSP’s robust detection performance - maintained under heterogeneous spikelet variability and electrode drift conditions - was preserved within the trained model and generalizes effectively to new, previously unseen datasets.

### 3.4 Ablation studies demonstrate the utility of the CUSP design components

CUSP outperformed all existing state-of-the-art spike sorting algorithms in our benchmarking dataset. To further understand how specific architectural and training components contributed to this high performance, we conducted a sequence of ablation studies to assess the contributions individual components on the learning dynamics and model accuracy during leave-one-out training and testing. Specifically, we performed three sets of ablations: 1) model structure ablation, 2) multi-channel input ablation, and 3) data augmentation ablation.

In the model structure ablation, we compared the CUSP full model with architecture variants, in which either the self-attention module or the U-net structure was removed. In the multi-channel input ablation, we systematically reduced CUSP’s spatial input scope from 12 channels to 8, 4, and finally a single-channel to evaluate the importance of multi-channel information. In the data augmentation ablation, the full training configuration to versions with individual augmentation techniques - channel drift, sample jitter, or voltage reversal - disabled. During each experiment, we tracked training and test iterations to assess convergence behavior and generalization.

Analysis of training and test loss traces from the model structure ablation revealed that the full CUSP model learned faster and achieved lower loss values than versions lacking either the self-attention component or U-net structure (**Fig. 6A**). These results indicate that both the self-attention and the U-net modules facilitate the learning of the latent CS feature representations during training, and improve evaluation performance during testing. In the multi-channel input ablation experiment, CUSP’s performance progressively degraded as the number of input channels was reduced, with higher training and test losses observed for fewer channels (**Fig. 6B**). This finding suggests that CUSP’s strong performance in complex spike sorting depends partly on its ability to integrate spatial information across multiple recording channels. Finally, ablations of the data augmentation strategies – including channel drift, sample jittering, and voltage reversal - also resulted in slower learning, and higher training and test loss values (**Fig. 6C**), demonstrating that these augmentation techniques can be beneficial for faster convergence and better generalization from the training to test dataset.

**Fig 6.**
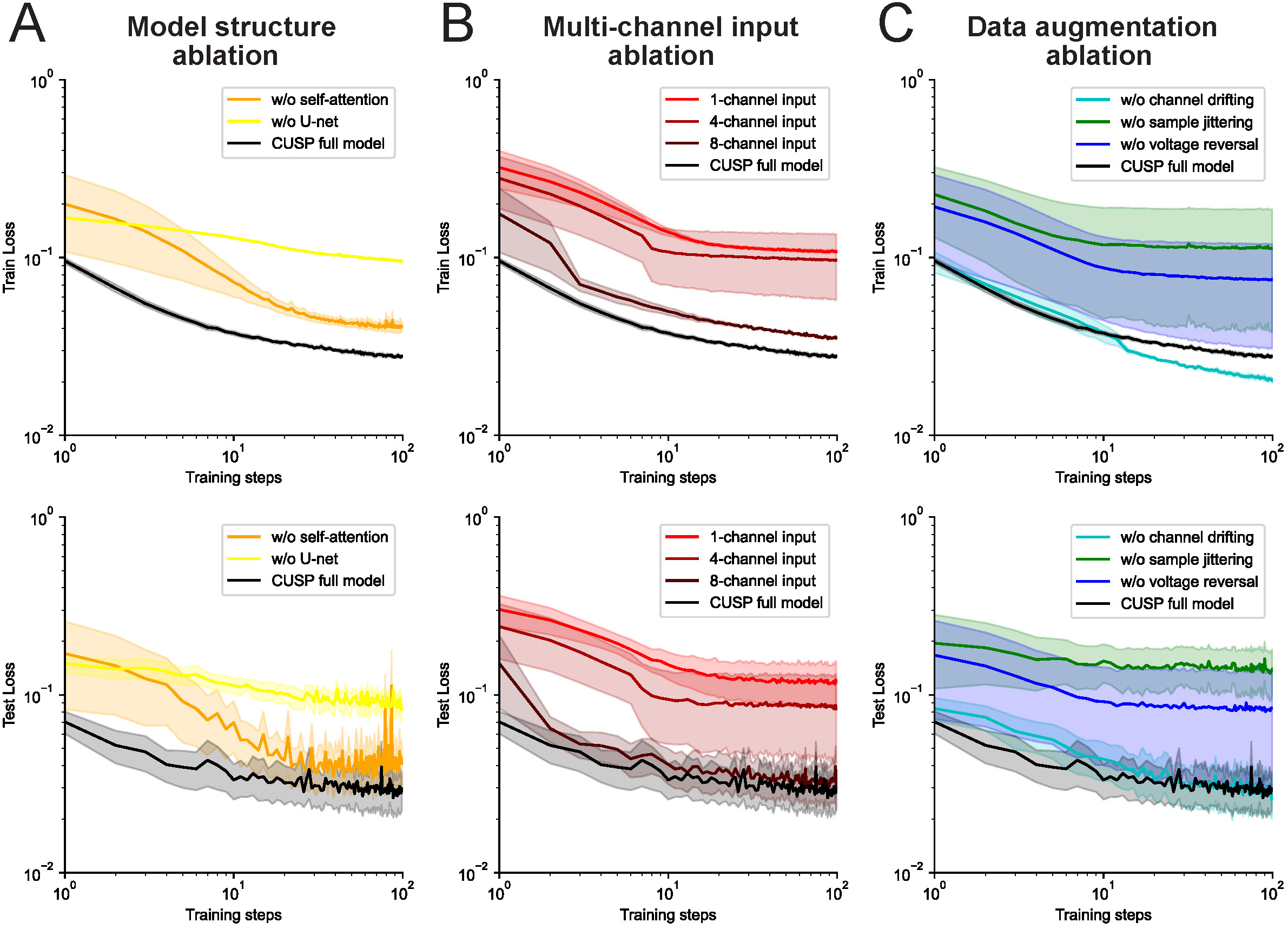
Ablation studies reveal utility of model designs in CUSP. **(A)** Averaged training and test loss traces over 100 training steps for CUSP full model (black), CUSP without self-attention components (orange), and CUSP without U-net structure (yellow) in the model structure ablation study. **(B)** The averaged training and test loss traces over 100 training steps for the CUSP full model with 12-channel input (black), CUSP with 8-channel input (dark red), CUSP with 4-channel input (middle red), and CUSP with only single-channel input (light red) in the multi-channel input ablation study. **(C)** The averaged training and test loss traces over 100 training steps for CUSP full model (black), CUSP without channel drifting (cyan), CUSP without sample jittering (green), and CUSP without voltage reversal (blue) in the data augmentation ablation study. The solid line and shadowed area indicate the mean ± standard error of the loss traces over training and testing.

We further benchmarked the ablation models on our CS dataset by using the same leave-one-out cross validation and compared their performance scores with those of the full CUSP model. Overall, the full CUSP model achieved the best CS sorting performance, exhibiting the highest precision (0.853±0.037), F1 score (0.834±0.032), and a leading recall (0.843±0.040) (**Table 1**). Model structure ablations that removed either the self-attention or U-net components resulted reduced overall performance, as reflected by lower F1 scores. A similar trend was observed in the multi-channel input ablations, where reducing the number of input channels led to a progressive decline in precision and F1 scores. Disabling data augmentation techniques, such as channel drift, sample jittering, or voltage reversal, also impaired model performance. Together, these results demonstrate that each architectural and training component contributes critically to CUSP’s performance, and that the full model is optimally designed for complex spike detection.

**Table 1.** Benchmarking performances in ablation studies.

| **Ablation settings** | **Precision** | **Recall** | **F1** |
| --- | --- | --- | --- |
| CUSP full model | **$0.853 \pm 0.037$** | $0.843 \pm 0.040$ | **$0.834 \pm 0.032$** |
| w/o self-attention | $0.783 \pm 0.059$ | $0.788 \pm 0.062$ | $0.776 \pm 0.057$ |
| w/o U-net | $0.810 \pm 0.044$ | **$0.879 \pm 0.034$** | $0.822 \pm 0.033$ |
| 1-channel input | $0.365 \pm 0.099$ | $0.849 \pm 0.044$ | $0.363 \pm 0.096$ |
| 4-channel input | $0.640 \pm 0.081$ | $0.780 \pm 0.076$ | $0.668 \pm 0.080$ |
| 8-channel input | $0.771 \pm 0.059$ | $0.806 \pm 0.062$ | $0.777 \pm 0.057$ |
| w/o channel drifting | $0.821 \pm 0.039$ | $0.803 \pm 0.052$ | $0.783 \pm 0.040$ |
| w/o sample jittering | $0.768 \pm 0.060$ | $0.790 \pm 0.061$ | $0.762 \pm 0.055$ |
| w/o voltage reversal | $0.766 \pm 0.060$ | $0.802 \pm 0.061$ | $0.770 \pm 0.055$ |

## 4. Discussion

In this study, we introduce a novel deep learning framework for automated sorting of complex spikes (CSs) from cerebellar Purkinje cells, a longstanding technical challenge in systems neuroscience. Analyzing CSs alongside simple spikes (SSs) is essential for understanding cerebellar computation, as these two classes of spikes convey complementary streams of information: CSs relay powerful instructive signals from climbing fibers, while SSs encode sensorimotor predictions. However, the unique morphology and rarity of CSs—combined with noisy recording environments, waveform variability, and electrode drift—have hindered efforts to develop automated and scalable sorting solutions. Our model addresses these challenges by integrating spatial and temporal information across high-density multichannel recordings using a U-net sequence-to-sequence architecture enhanced with hybrid self-attention inception blocks. This design enables precise CS onset detection while remaining robust to variability, achieving expert-level performance in both controlled and realistic conditions. Importantly, this same robustness and scalability make CUSP a more general tool that can be invariant to dynamically complex spike patterns that are known to be poorly captured by the more prevalent simple spike based sorting algorithms. Such an open tool could provide a novel framework for studying information coding across circuits.

We benchmarked CUSP against leading single-channel and multi-channel spike sorting algorithms, demonstrating substantial improvements in detection accuracy, particularly under challenging recording conditions (**Fig. 4**). Notably, our method preserved performance when tested with real-world datasets characterized by waveform distortion, spikelet variability, and low signal-to-noise ratios (SNRs)—conditions that frequently diminish the reliability of other algorithms. These results underscore the utility of our approach for cerebellar recordings at scale, where manual annotation becomes impractical and conventional methods often fail. A key innovation is the model’s end-to-end utilization of multi-channel data. While prior CS sorting methods have predominantly relied on single-channel waveforms (Zur & Joshua, 2019; Markanday et al., 2020; Sedaghat-Nejad et al., 2021), our model processes the full spatial context of MEA recordings. For instance, Zur & Joshua (2019) demonstrated that coupling between SS and CS waveforms can aid classification by analyzing low-frequency features via PCA. However, this approach—restricted to single-channel recordings—struggles to generalize across recordings due to variability in CS waveform expression; many CSs lack prominent low-frequency signatures, limiting detection reliability. A recent strategy that used deep learning to classify CSs in single-channel recordings based on high-SNR was able to achieve good performance under optimal conditions (Markanday et al., 2020). P-sort (Sedaghat-Nejad et al. 2021) presents the Markanday et al. classifier in a more user-friendly interface with enhanced visualization tools using PCA and/or UMAP. While these additions offer greater flexibility, existing methods remain limited by their reliance on subjective manual input and their failure to leverage essential spatial information, which restricts scalability for high-density MEA datasets.

As a consequence, they cannot take advantage of spatial information critical for tracking CS activity across electrodes and become susceptible to noise, electrode drift, and overlapping signals—factors that are prevalent in naturalistic recordings.

By contrast, CUSP processes both local field potential (LFP) and action potential (AP) traces from multiple nearby channels, extracting robust spatial signatures that distinguish true CSs from background activity. The inclusion of self-attention layers enables adaptive weighting of relevant features, that can enhance generalization across animals, sessions, and electrode geometries. Indeed, the architectural structure of our approach yields strong performance across a variety of test conditions. Importantly, the ability to learn generalized spatiotemporal features directly from data obviates the need for handcrafted feature selection or template matching. This challenge parallels recent work in behavioral neuroscience, where even highly stereotyped motor patterns—obvious to human observers—require tailored computational tools for reliable detection from keypoint data. For example, Stanley et al. (2024) recently illustrated this in their analysis of circling behavior in mice.

To better understand how these architectural advantages translate into performance gains, we analyzed detection discrepancies between our model, expert annotators, and competing algorithms (**Fig. 5C**). Although many CSs detected exclusively by our model displayed clear, intact waveforms, they were nonetheless overlooked by expert annotators, who also showed considerable inter-rater variability. In contrast, other algorithms showed distinctive error profiles: for example, Kilosort4 produced misaligned false positives due to erroneous spikelet clustering, while Spyking Circus2 failed to detect a large portion of high-confidence CSs under low-SNR conditions, which is not surprising given the propensity of existing algorithms to over-split even SS events due to minor waveform changes (Koukuntla et al. 2025). These results underscore the advantage of our model’s ability to balance sensitivity and specificity across a wide range of recording environments.

Beyond detection robustness, CUSP addresses another critical barrier in the field: intra-cellular variability in CS morphology. It is well established that the fine temporal structure of CS waveforms—including the number, timing, and amplitude of spikelets—can vary substantially even within the same Purkinje cell (Yang & Lisberger, 2014; Lang et al., 2014; Warnaar et al., 2015; Zang et al., 2018; Burroughs et al., 2017). This variability is not mere noise: it has long been recognized that the precise spatiotemporal patterning of complex spike trains shapes Purkinje cell simple spike output, thereby dynamically linking climbing fiber input to plasticity and cerebellar computation (Llinás & Sugimori, 1980; Davie et al., 2008; Najafi & Medina, 2013; Zang et al., 2018; Streng et al., 2018). Consequently, tools that misclassify subtle waveform variants risk erasing biologically meaningful signals (Koukuntla et al. 2025).

CUSP, by design, accommodates this variability. Rather than enforcing rigid templates or clustering in low-dimensional PCA space, it leverages multiscale spatiotemporal features to consistently detect CSs with varying spikelet compositions within the same cell (**Fig. 5A**), enabling downstream analyses of how these patterns relate to motor learning or behavioral state. This capability not only advances spike sorting but also opens new opportunities for studying how information is encoded within CS morphology—work previously constrained by labor-intensive manual labeling (Markanday et al., 2019). Moreover, the same framework may extend beyond the cerebellum to other systems where neurons generate burst-like or dynamically complex activity distinct from their primary spikes. For example, hippocampal pyramidal neurons produce complex spike bursts that play critical roles in computation and plasticity yet remain difficult to isolate with standard tools (Grienberger et al., 2014; Raus Balind et al., 2019; Sneider et al., 2006; Gao et al., 2021). By accommodating waveform variability, CUSP offers a generalizable approach for detecting such events, enabling new insights into their function across circuits.

Importantly, our method also performs well in situations with highly non-stationary recording conditions. As MEA recordings increase in duration and density, electrode drift and changes in background activity present major obstacles for both manual curation and automated tools (Windolf et al., 2025). The performance of many existing methods—including those using clustering or static filters—degrades sharply in such conditions. Our architecture’s use of convolutional layers combined with self-attention blocks allows it to continuously re-weight salient features, mitigating the impact of drift or spatial nonuniformity. This flexibility enables our model to maintain high accuracy without needing retraining or user intervention mid-session, making it particularly well suited for large-scale, longitudinal recordings.

From a broader perspective, our results support a growing shift toward data-driven, end-to-end learning models in the field of systems neuroscience (Vogt 2018, Xu et al. 2023, Chen et al. 2025). While template-matching and clustering-based methods have historically dominated spike sorting (e.g., Pachitariu et al. 2024, Chung et al., 2017), these techniques are increasingly limited by their reliance on human tuning, narrow assumptions about spike shape, and limited scalability. The use of deep neural networks—particularly those capable of learning from multi-channel LFP and AP signals transformed into analytic representations via the Hilbert transform —represents a powerful alternative. Crucially, our approach does not require retraining for each recording or experiment. Once trained, it generalizes robustly across datasets, offering immediate utility to laboratories using MEA systems to study cerebellar function. To facilitate future development and comparative benchmarking, we have also curated a large, open-access dataset comprising diverse cerebellar recordings (https://github.com/CullenLab/CUSP). This resource includes expert annotations, low-and high-SNR examples, and challenging segments with waveform variability or drift.

The dataset, alongside our trained model and evaluation code, provides a standardized framework for evaluating future methods. By establishing clear performance baselines and highlighting common failure modes, it can accelerate progress toward more general and interpretable spike sorting solutions.

Despite the fact that data-driven machine learning methods are often viewed as lacking interpretability, CUSP affords several interpretable mechanistic insights within its model architecture. The model’s design and signal processing paths are designed to capture distinct aspects of LFP and AP inputs: the broad, wide-range complex spike envelope features and the fine temporal structure of spike and spikelet patterns. This dual-path architecture likely explains the model’s high detection accuracy for CSs with spikelet diversity. Moreover, the inception block’s multi-scale filters enable ablation-based attributions by systematically quantifying the contributions of individual convolutional kernel branches operating at different temporal scales. The U-net structure further contributes to performance by using skip connections that bridge early-stage, low-level temporal and spatial features from the encoder with the late-stage, high-level, abstract representations in the decoder – an arrangement that enhances precise CS onset detection. Finally, the incorporation of self-attention mechanisms within the inception blocks yields self-adaptive spatiotemporal weight maps, allowing the network to automatically identify which channel or time segments most strongly influence CS detection. This flexibility across both channel and sample provides robust performance, even under conditions involving sample jittering and channel drift.

Although CUSP achieved high performance in CS detection, it remains a supervised learning algorithm, and therefore bears several potential systematic limitations.

Supervised CS detection benefits from expert-provided ground-truth labels, which encode domain-specific knowledge about the CS waveform features such as spikelet diversity. However, these labels also implicitly capture high-level background information, including the estimated position of Purkinje cells in the cerebellar cortex layer structure, the electrode trajectory, and functional connectivity among neighboring cell type that help predict Purkinje cell identity. While such supervised knowledge embedded in the CS labels enhances performance, it is also labor-intensive to obtain due to the time-consuming nature of manual expert curation. Moreover, individual biases among human experts are inevitably transmitted through their annotations, leading to a supervised model whose performance cannot exceed the upper bound set by human-level detection accuracy.

In this study, CUSP was trained and benchmarked against human-level CS detection using consensus-labeled datasets derived from multiple experts to minimize individual bias during both training and evaluation. Nevertheless, purely supervised learning remains constrained by the limited availability and subjectivity of expert labels. In contrast, unsupervised or self-supervised learning approaches - such as contrastive learning or autoencoder based representation learning - could leverage the large volume of unlabeled CS data to extract robust and generalizable CS features. However, unsupervised clustering alone, as commonly implemented in existing spike sorters, is insufficient for CS detection due to the spikelet variability across CS as well as their relative rarity. Taken together, future improvement in CS detection model could benefit from semi-supervision strategies. One promising direction would be to pretrain the encoder within an autoencoder-like architecture using large unlabeled datasets to learn robust latent representations, followed by fine-tuning on a modest number of expert-labeled CS examples - a few-shot learning framework (Fang & Zamani, 2025). Another complementary approach could employ a crowdsourcing strategy that integrates features extracted from multiple unsupervised spike-sorting algorithms and then trains a supervised model to identify CS from this ensemble-derived feature space (Wyngaard et al., 2022).

Thus, in summary, our work advances the field in several key respects. First, it introduces a high-performing deep learning model that detects CSs with high precision and robustness, even under noisy and nonstationary conditions. Second, it explicitly incorporates multichannel spatial features, improving generalization beyond traditional single-channel methods. Third, it accommodates biologically meaningful CS waveform variability, enabling future studies of information coding and plasticity in the cerebellum. Finally, it provides an open benchmarking resource that supports ongoing methodological innovation. Together, these contributions represent a significant step forward in cerebellar systems neuroscience, bridging the gap between manual expertise and scalable automation. Moreover, by combining robustness to waveform variability and electrode drift with scalability across high-density recordings, CUSP establishes a broadly applicable framework for detecting complex and/or burst-like neural events beyond the cerebellum, opening new opportunities for studying information coding across diverse circuits.

## Acknowledgement

We thank Dr. M. Hu for his assistance in manual annotations of complex spikes from datasets. This work was supported by funding from the National Institute on Deafness and Other Communication Disorders (R01-DC002390 and R01-DC018061) (K.E.C. and A.S.C) Raynor Cerebellar Project (K.E.C. and A.S.C), as well as a Natural Sciences and Engineering Postdoctoral Fellowship and Kavli Neuroscience Discovery Institute Distinguished Postdoctoral Fellowship (R.M.).

## Author contributions

C.B. designed and coded the algorithms, prepared datasets, analyzed results, prepared figures, and drafted the manuscript.

R.M. conducted the experiments, prepared the datasets, prepared figures, and provided revisions to the manuscript.

A.S.C. provided guidance on the algorithms and revisions to the manuscript.

K.E.C. designed experiments, supervised the project, and wrote the manuscript.

## Ethical statements

All experimental procedures were conducted with approval from the Johns Hopkins University Animal Care and Use Committee and adhered to the guidelines outlined by the United States National Institutes of Health (Protocol PR22M342). The datasets used in this paper is prepared from cerebellar recordings performed on one female and one male macaque monkeys (Macaca mulatta), weighing 7 and 12kg, respectively. A total of 63 recording sessions were performed. Throughout the course of the study, these animals were maintained in a controlled environment with a 12-hour light/dark cycle. Both macaque monkeys had previously participated in other research studies within our laboratory, exhibited overall good health, and did not require any medication throughout the course of this experiment.

---

**Fig S1.**
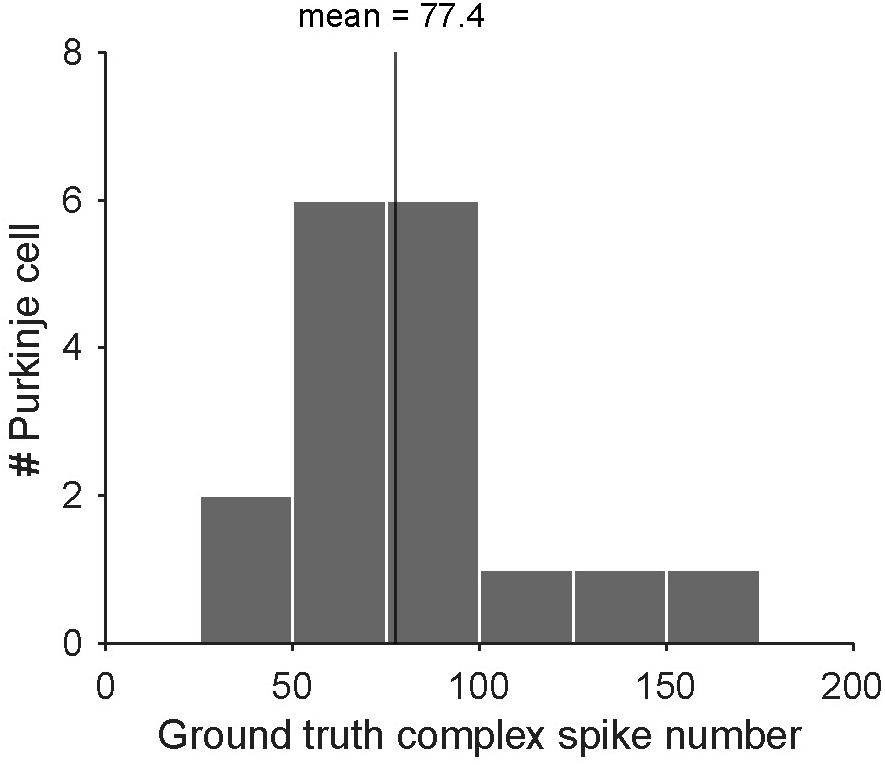
Histogram of ground truth expert labeled complex spike number per number in our dataset. In average, leaving out 1 Purkinje cells will exclude 77.4 ground truth expert labeled CS event instances from training.

**Fig S2.**
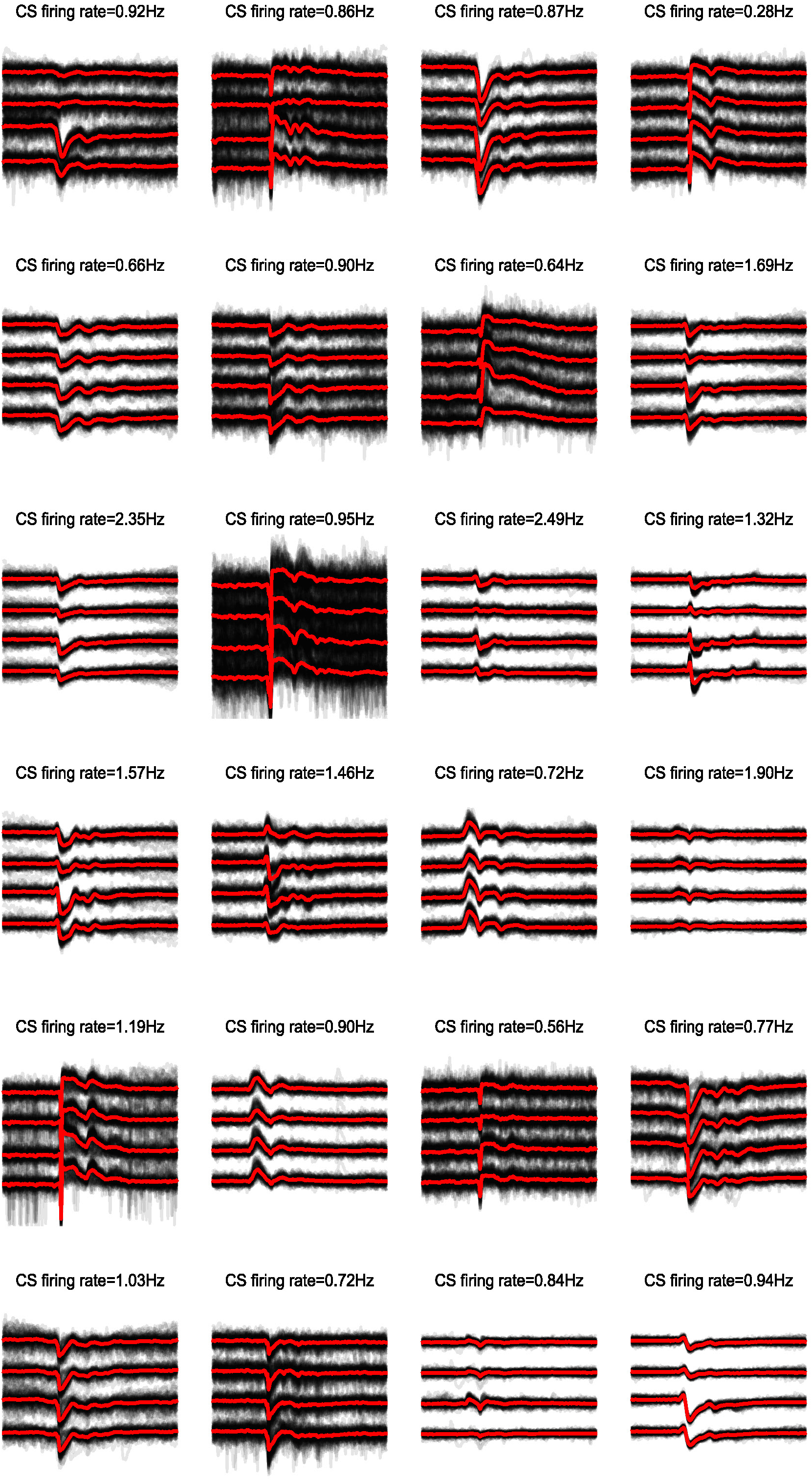
CUSP-detected complex spike examples from recording database outside the reported expert-labeled complex spike datasets. Each complex spike example contains four-channel waveforms adjacent to the complex spike peak channel (black), the averaged waveform (red), and its averaged firing rate at resting discharge.

**Fig S3.**
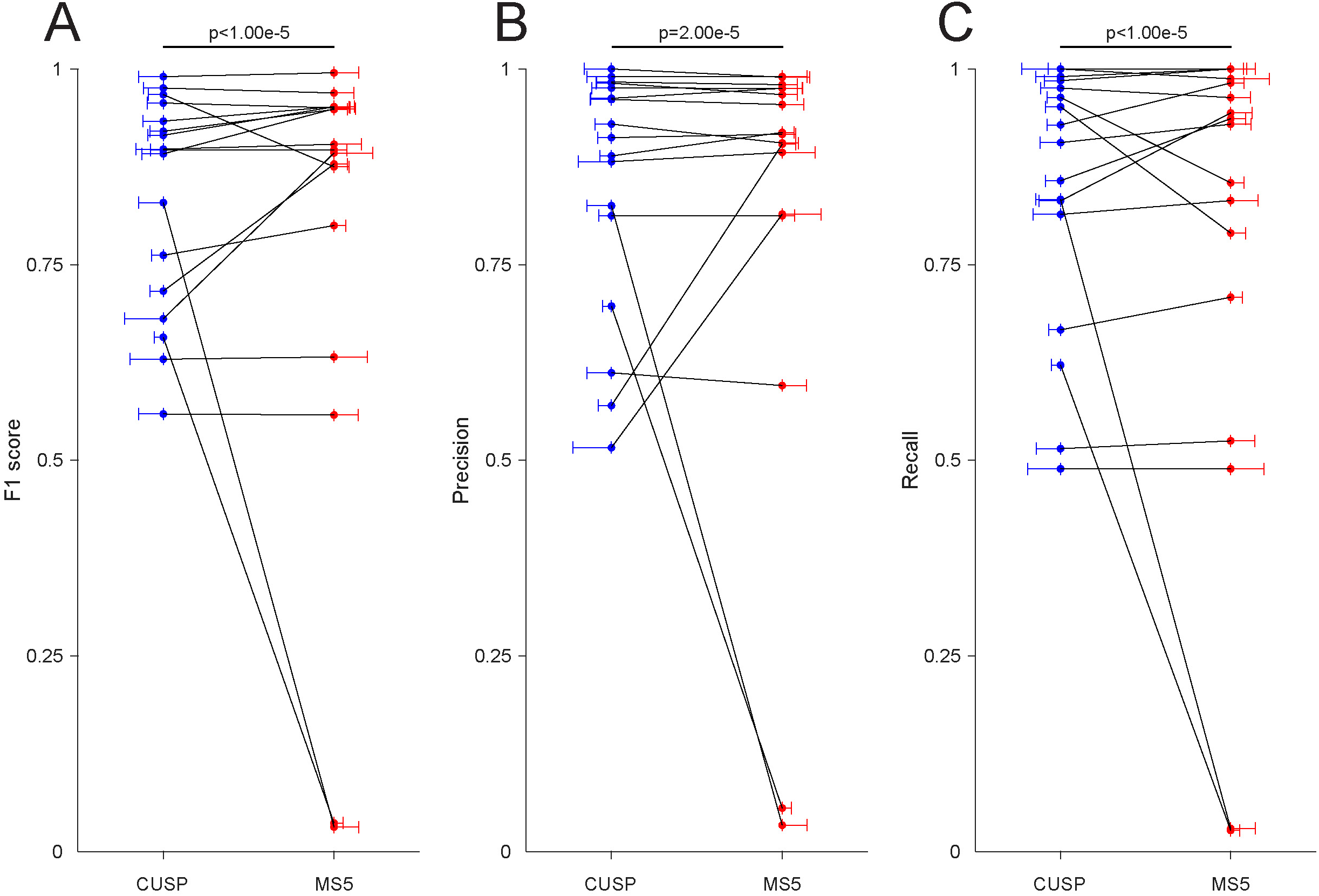
CUSP shows significantly higher complex spike sorting performance than the existing state-of-the-art spike sorter. Paired permutation tests were performed between CUSP and the existing state-of-the-art spike sorter (Mountainsort5) to compare **(A)** F1 scores, **(B)** precision scores, and **(C)** recall scores during the leave-one-out cross validation. The randomized permutations were re-weighted based on the total expert labeled complex spike numbers in each session to compensate for the imbalance among the variable session time lengths and complex spike firing rates and then repeated for 100,000 times. Each black line is one session and each dot is the score value of CUSP (blue) or Mountainsort5 (red). The length of the horizontal bar near each dot is proportion to the permutation weight (total expert labeled complex spike number) in that session. F1 score: p<1.00e-5; precision score: p=2.00e-5; recall score: p<1.00e-5.

**Fig S4.**
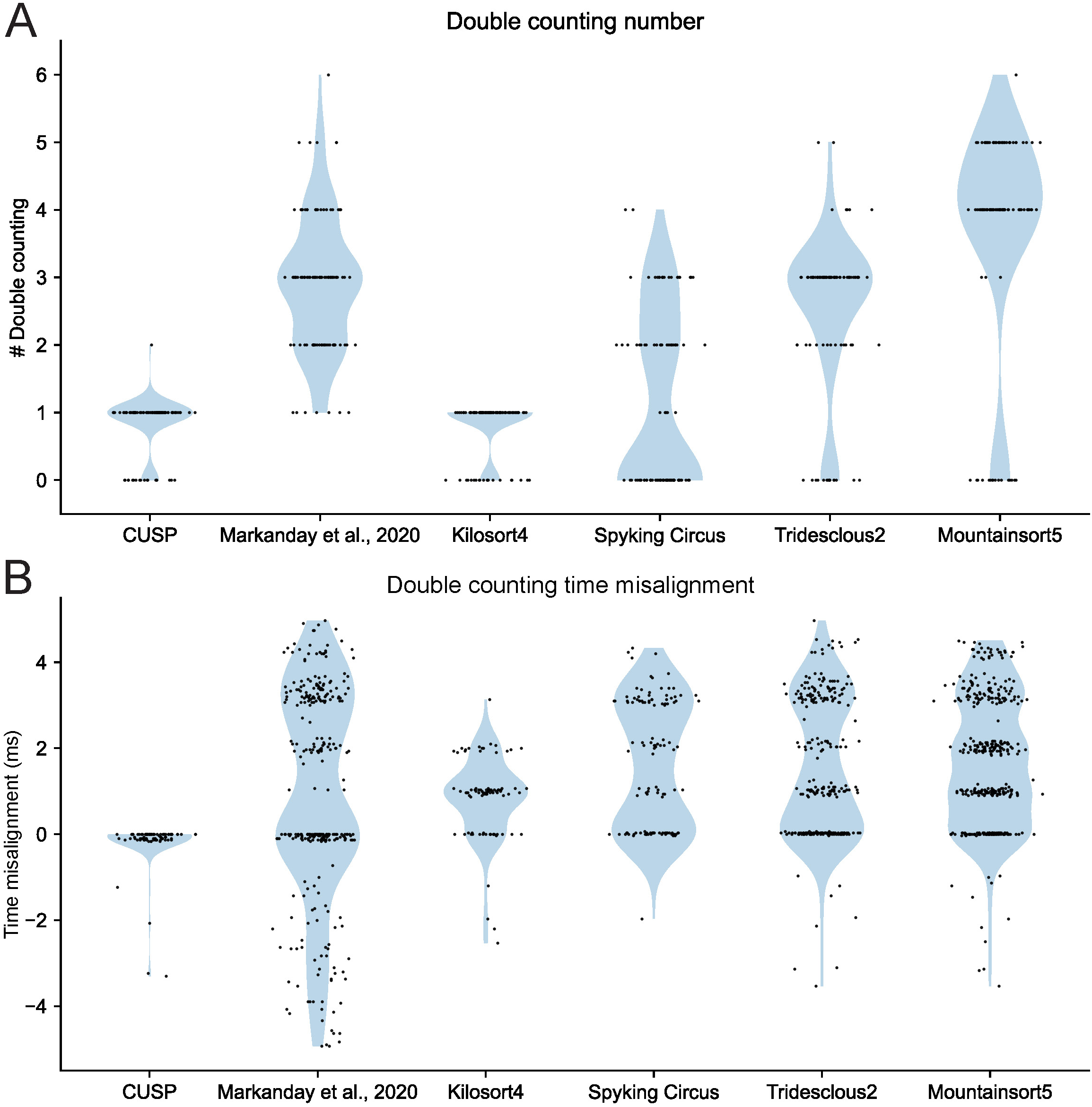
CUSP shows less double-counting problems when sorting complex spikes with spikelet diversity. **(A)** Double-counting number counting for each algorithm in an example session when sorting complex spikes with variable spikelet numbers and inter-spikelet intervals. Each black dot is the multiple-counting number of each algorithm detections for each expert-labeled complex spike. The blue area shows the general double counting number distributions for each algorithm. **(B)** Double-counting time misalignment for each algorithm in the same example session. Each black dot is the time misalignment of each algorithm detection relative to the expert-labeled complex spike onset time. The blue area shows the general double counting time misalignment distributions for each algorithm.

**Fig S5.**
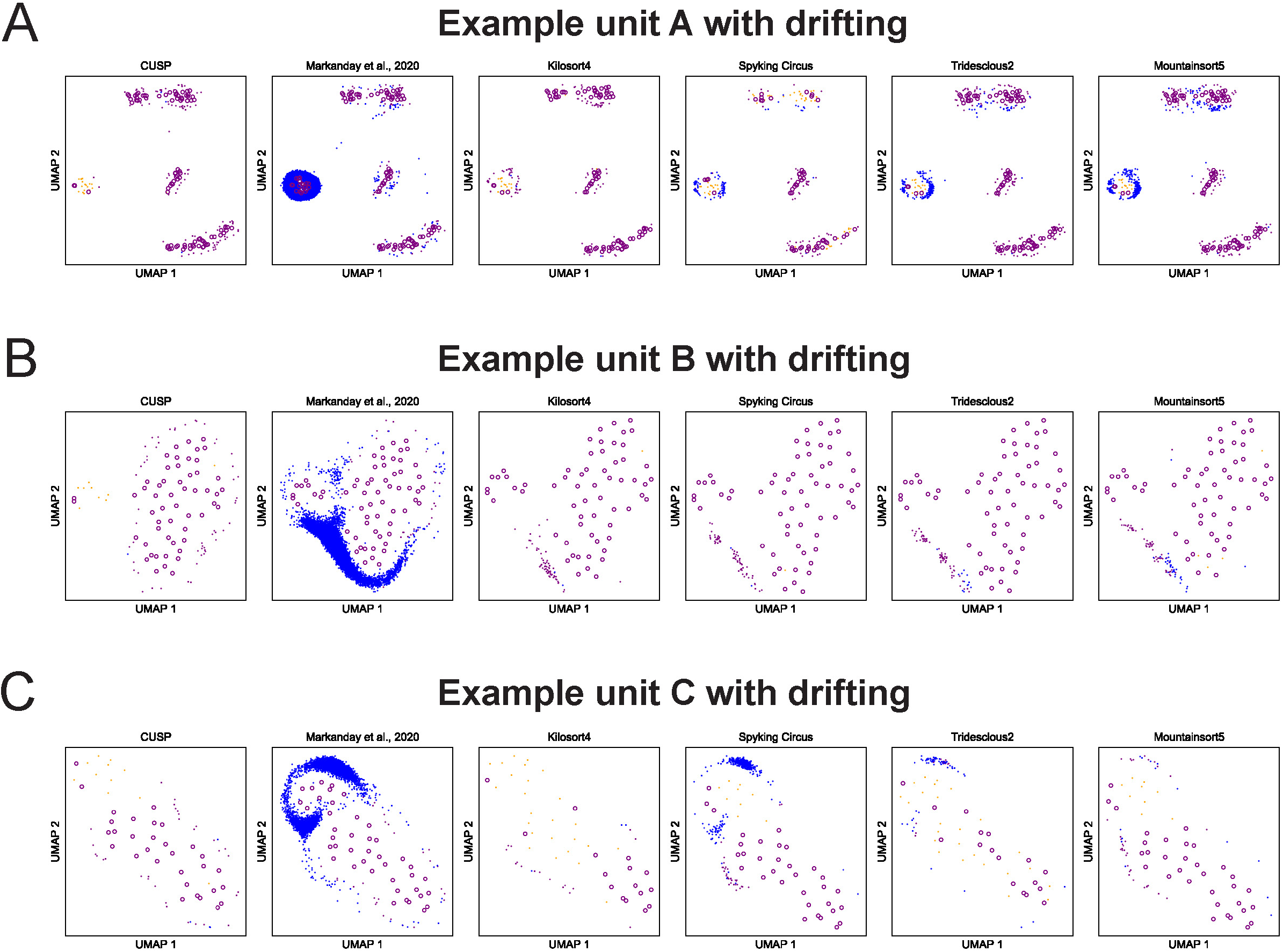
CUSP shows its detected complex spike cluster distributions aligned with the manifold of the expert labeled complex spikes in the embedded space. **(A-C)** The expert-labeled complex spike waveforms from three example units with recording drifting were embedded into two-dimensional manifold using UMAP, as well as with each algorithm detected complex spike waveforms projected into the same two-dimensional space using UMAP. The purple circle indicates each successfully detected expert-labeled complex spike; the purple dot indicates each algorithm detected complex spike that was matched to an expert-labeled complex spike; the blue dot indicates complex spike labeled only by each algorithm; the yellow dot indicates the complex spike labeled only by experts.

